# SMRT-Cappable-seq reveals complex operon variants in bacteria

**DOI:** 10.1101/262964

**Authors:** Bo Yan, Matthew Boitano, Tyson Clark, Laurence Ettwiller

## Abstract

Current methods for genome-wide analysis of gene expression requires shredding original transcripts into small fragments for short-read sequencing. In bacteria, the resulting fragmented information hides operon complexity. Additionally, *in-vivo* processing of transcripts confounds the accurate identification of the 5’ and 3’ ends of operons. Here we developed a novel methodology called SMRT-Cappable-seq that combines the isolation of unfragmented primary transcripts with single-molecule long read sequencing. Applied to *E. coli*, this technology results in an unprecedented definition of the transcriptome with 34% of the known operons being extended by at least one gene. Furthermore, 40% of transcription termination sites have read-through that alters the gene content of the operons. As a result, most of the bacterial genes are present in multiple operon variants reminiscent of eukaryotic splicing. By providing an unprecedented granularity in the operon structure, this study represents an important resource for the study of prokaryotic gene network and regulation.

## Introduction

The model of operon structure proposed by Jacob and Monod invoked a group of contiguous genes regulated by a common promoter in bacteria [1]. The resulting polycistronic transcripts can be several kilobases (kb) long and encode several proteins usually associated with the same function or metabolic pathway.

With decades of research on bacterial gene regulation and the advance of high throughput technologies, this operon model has been found to be complex with a rich landscape of gene regulation mechanisms. For examples, riboswitches sense endogenous regulatory ligands and upon ligand binding adopt either a terminator or an antiterminator structure [2, 3]. Riboswitches have been found in the 5’ leader regions generating either aborted transcripts or full-length transcripts [2, 3]. Furthermore, genome-wide studies have identified more transcription start sites (TSS) and transcription termination sites (TTS) than known operons [4-6], which hints at a more complex operon structure than previous thought.

For genomewide TSS determination, we and others have used the fact that the original 5’ end of prokaryotic primary transcripts are triphosphorylated, while processed 5’ ends are not and device methodologies to sequence specifically the TSS at base resolution [4, 7]. For genomewide TTS determination, such molecular distinction is absent and previous methods, such as term-seq, rely on the sequencing of original and processed 3’ ends of RNA in bacteria [8]. All experimental methods for the identification of such transcriptional landmarks in bacteria rely on the necessary fragmentation of transcripts for short read sequencing. Consequently, the larger context by which genes are expressed and the phasing between starts and the ends of primary transcripts are lost.

Yet the ability to simultaneously sequence both TSS and TTS of primary transcripts at single molecule resolution is key to the accurate identification of operons. Sequencing full-length cDNA using long read sequencing technology has been previously described in Eukaryotes (iso-seq) [9, 10] and has revealed a complex array of isoforms [11], but so far no similar technologies has been developed for prokaryotes.

### SMRT-Cappable-seq

In this study we have developed a novel strategy combining the isolation of the full-length prokaryotic primary transcriptome with PacBio SMRT (Single Molecule, Real-Time) sequencing. To ensure the sequencing of full-length primary transcripts, we capture the triphosphorylated 5’ ends of transcripts matching the TSS. As the first step of the *in-vivo* RNA degradation pathway is thought to consist of the removal of the 5’ triphosphate [12], the capturing of the 5’ triphosphate should also ensure the removal of degraded and/or processed transcripts. While indirect, this strategy is expected to markedly enriched for primary transcripts that have also retained their original TTS.

For Long read sequencing, we adapted Cappable-seq technology previously used to identify TSS [4] for the isolation of long (>1kb) transcripts (**Figure 1A, Figure S7**). The principle of the Cappable technology is based on specific desthio-biotinylation of the 5’ triphosphate characteristic of the first nucleotide incorporated by the RNA polymerases [4]. The desthio-biotinylated RNA is captured on streptavidin beads and subsequently released from the beads after several washing steps to remove processed RNA. To capture the 3’ end of the transcripts, a poly A tail is added to the 3’ end of RNA and cDNA is synthesized via reverse transcription (RT) using an anchored poly dT primer. After RT reaction, a polyG linker is added to the 3’ end of cDNA for second strand synthesis using Terminal transferase. Finally, the unfragmented cDNA is size selected for large fragments (>1kb), amplified and sequenced using PacBio long read sequencing technology, resulting in the identification of full-length transcripts at base resolution. Importantly, long read sequencing provides the phasing (pairing) of both ends of transcripts, overcoming the inabilities of short reads to obtain long-range continuity (Figure 1D). Thus, SMRT-Cappable-seq provides for the first time a powerful approach for directly identify entire operon at molecule resolution in bacteria.

**Figure 1:**
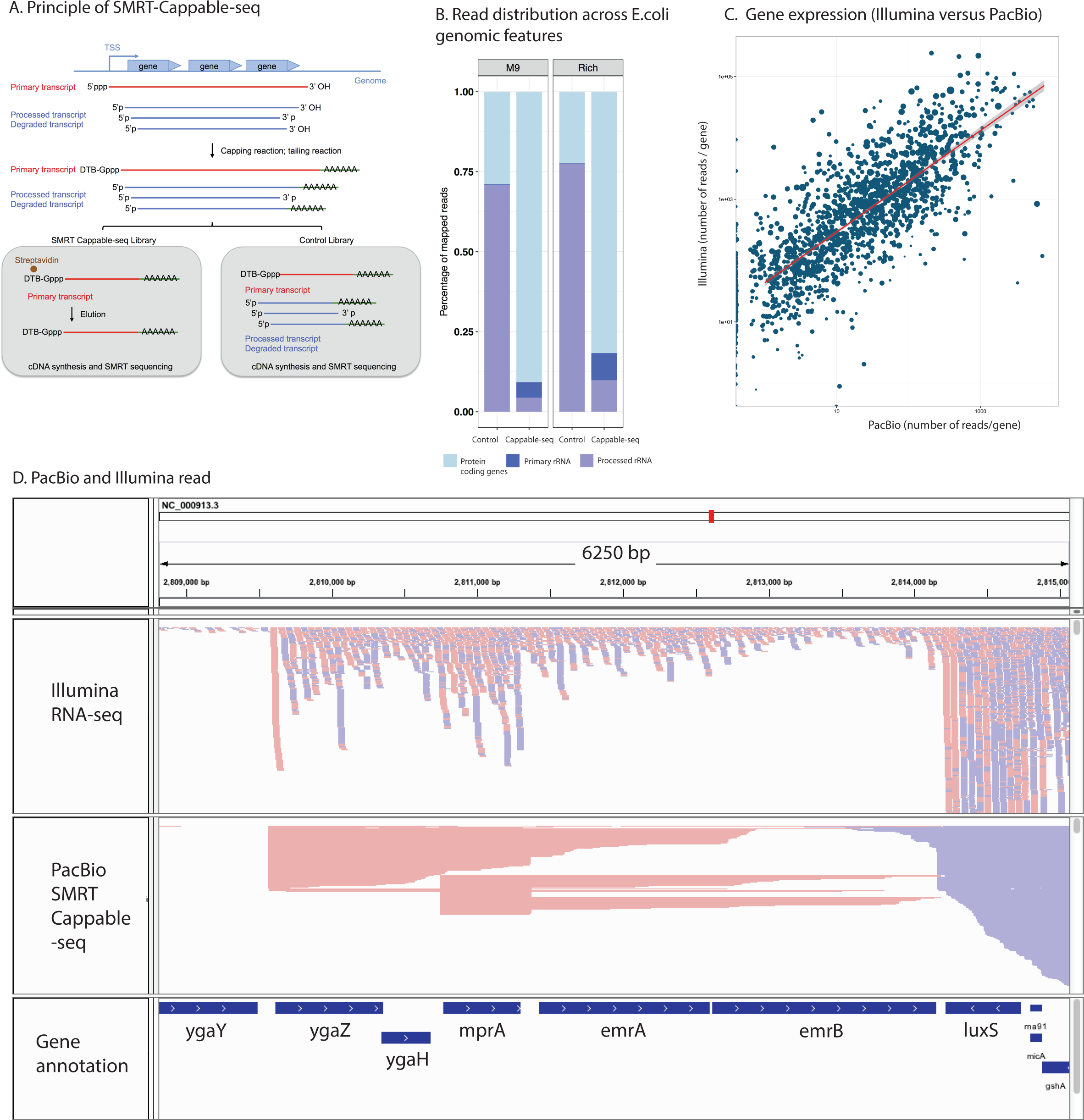
SMRT-Cappable-seq identifies full-length transcripts in bacteria. **A**. Schema of the SMRT-Cappable-seq methodology. 5’ triphosphorylated transcripts are capped with a desthio-biotinylated cap analog and bound to the streptavidin beads to specifically capture primary transcripts starting at TSS. The polyadenylation step ensures the priming of the anchored poly dT primer for cDNA synthesis at the most 3’ end of the transcript. **B**. Percentage of of reads mapping to protein coding genes (light blue), primary rRNA (dark blue) and processed rRNA (purple) for both M9 and Rich growth conditions (left panel and right panel respectively). Reads mapped to primary rRNA are defined as reads which start at a known TSS of a primary rRNA transcript [4], while processed rRNAs correspond to reads mapped to the rRNA genes but do not start at these TSSs. SMRT-Cappable-seq libraries show enrichment of primary transcripts (primary rRNA + protein coding genes relative to rRNA) compared to control libraries. C. Comparison between gene expression level (total number of reads mapping to annotated genes) for illumina RNA-seq and SMRT-Cappable seq. Point size denotes the size of the gene (bp). D. Integrative Genomics Viewer (IGV) representation of the mapping of SMRT-Cappable-seq reads (top) compared to Illumina RNA-seq reads (bottom) in the mprA locus. Forward oriented reads are labeled in pink, reverse oriented reads are label in blue.

We applied SMRT-Cappable-seq to unfragmented total RNA from *E. coli* grown in minimal (M9) or Rich medium to compare how growth conditions affect the full-length transcriptome. We combined both data sets to obtain a comprehensive overview of the transcriptional landscape of long transcripts (>1kb) in *E. coli.* The combined dataset consists of a total of half a million sequencing reads with an average length of 2000 bp. More than 99% of reads are mapped to the *E. coli* genome and only 0.2% of the reads have chimera structures (**Material and Methods**). With a coverage of 90.3 % of the genome and 81% of the known genes covered entirely by at least one read, SMRT-Cappable-seq provides a comprehensive view of the bacteria full-length transcriptome. Despite size selection of the SMRT-Cappable-seq library, we found a decent correlation (corr = 0.46) between gene expression derived from SMRT-Cappable-seq and published Illumina RNA-seq [13] (**Figure 1 C and D and Supplementary text**) indicating that SMRT-Cappable-seq is suitable for quantitative measurement of transcript levels.

To assess the depletion of processed transcripts, which is an important step in the SMRT-Cappable-seq procedure, we analyzed the fraction of reads that are mapped to ribosomal RNA (rRNA). In bacteria, rRNAs are formed by the processing of a single pre-rRNA transcript to form the mature 5S, 16S and 23S rRNAs [14]. Because processed rRNAs accounts for the vast majority of the RNA in the cell [14], the differential levels of processed rRNA in the control versus SMRT-Cappable-seq libraries is a good indicator of the specificity of SMRT-Cappable-seq for primary transcripts. We found that the reads from processed rRNA drops from 71% in the control library to 4% in the SMRT-Cappable-seq library (**Figure 1B and Supplementary text**). Furthermore, amongst the reads mapped to rRNA, only 0.4% of rRNA reads in the control library corresponds to pre-rRNA transcript (based on the 5’ end location) versus up to 53% in the SMRT-Cappable-seq library (**Supplementary text**). Accordingly, the fraction of mappable reads matching known 5’ or 3’ processing sites is only 2% in the SMRT-Cappable-seq libraries, sharply contrasting with 33% in the control library (**Material and Methods, Supplementary text**).

These results are validated by qPCR experiments designed to measure the absolute recovery of primary and processed transcripts, before and after streptavidin enrichment. By measuring the absolute amount of RNA molecules, qPCR provides a more accurate estimation of the enrichment for primary transcript compared to sequencing. qPCR results reveal that around 30% of primary transcripts are recovered after streptavidin enrichment compared to only 0.01% recovery rate for processed ribosomal RNAs (**Figure S1**). Based on these qPCR results, we conclude that SMRT-Cappable-seq has a 1000-fold greater recovery of primary transcripts compared to processed RNAs. Taken together, our results demonstrate that SMRT-Cappable-seq enriches for primary transcripts and avoids sequencing of processed transcripts such as ribosomal RNA, tRNA and others.

### Transcription start sites

Consistent with the enrichment for primary transcript, we found that 93% of mapped SMRT-Cappable-seq reads start within 5 bp of a previously annotated TSS [4] compared to only 15% in the control library (**Supplementary Text**). We define TSS at single base resolution as genomic positions containing a significant accumulation of 5’ end reads. If several TSSs were found within a 5 bp window, only the TSS position with the highest score was retained (**Material and methods**). Libraries derived from M9 and Rich growth conditions have a total of 2186 and 1902 TSSs, respectively. Comparison between growth conditions reveals that 1350 TSSs are common and the TSS usage is globally correlated (corr =0.3). The prefered nucleotide for TSS is either adenosine or guanosine consistent with the litterature [15] and nucleotide bias at −10 and −35 bp reveals the standard profile of bacterial promoters (**Figure S2A**). Collectively, these results confirm the 5’end completeness of the majority of transcripts in SMRT-Cappable-seq.

### Transcription termination sites

Similar to the TSS, TTSs were define at single base resolution as genomic positions containing a significant accumulation of 3’ end reads. Contrasting with the 5’ end for which the labelling of the triphosphate ensures the direct identification of TSS, the capturing of the TTS is indirect and therefore requires more stringent filtering criteria (**Material and methods**). Consequently, we observed a lower fraction of transcripts containing TTS compared to TSS. Thus, the prediction of TTS results in 347 highly confident TTS in M9 condition, among which 74 and 1 have been previously identified as Rho independent and Rho dependent termination sites respectively. Very similar results were obtained for the Rich condition (**Supplementary text**). We compared our defined termination sites with the RnaselII cleavage sites, a post-transcriptional step that generates processed 3’ ends [16], and only 7 SMRT-Cappable-seq TTS sites are overlapping with RnaselII cleavage sites (**Supplementary text**). These results indicate that a minimal number of these TTSs are due to processing. Conversely, analysis of the sequence upstream of termination sites identifies two GC rich regions at −24 to −20 bp and −10 to −7 bp followed by a run of uracil residues characteristic of the Rho independent TTS (**Figure S2B**) [17].

The majority of termination sites defined by SMRT-Cappable-seq are located in intergenic regions (86%). Almost all of the TTS are located downstream of one or multiple gene(s), positioning these termination sites at the 3’ UTR ends of mature transcripts (See the example shown in **Figure 2A**). The relatively low abundance of premature termination sites found in the 5’ UTR region of genes can be explained by the size selection step during library preparation favoring longer transcripts. We nonetheless identify well known examples of termination sites in the 5’ UTR region of the operon such as the Leu operon (**Figure S3**).

**Figure 2:**
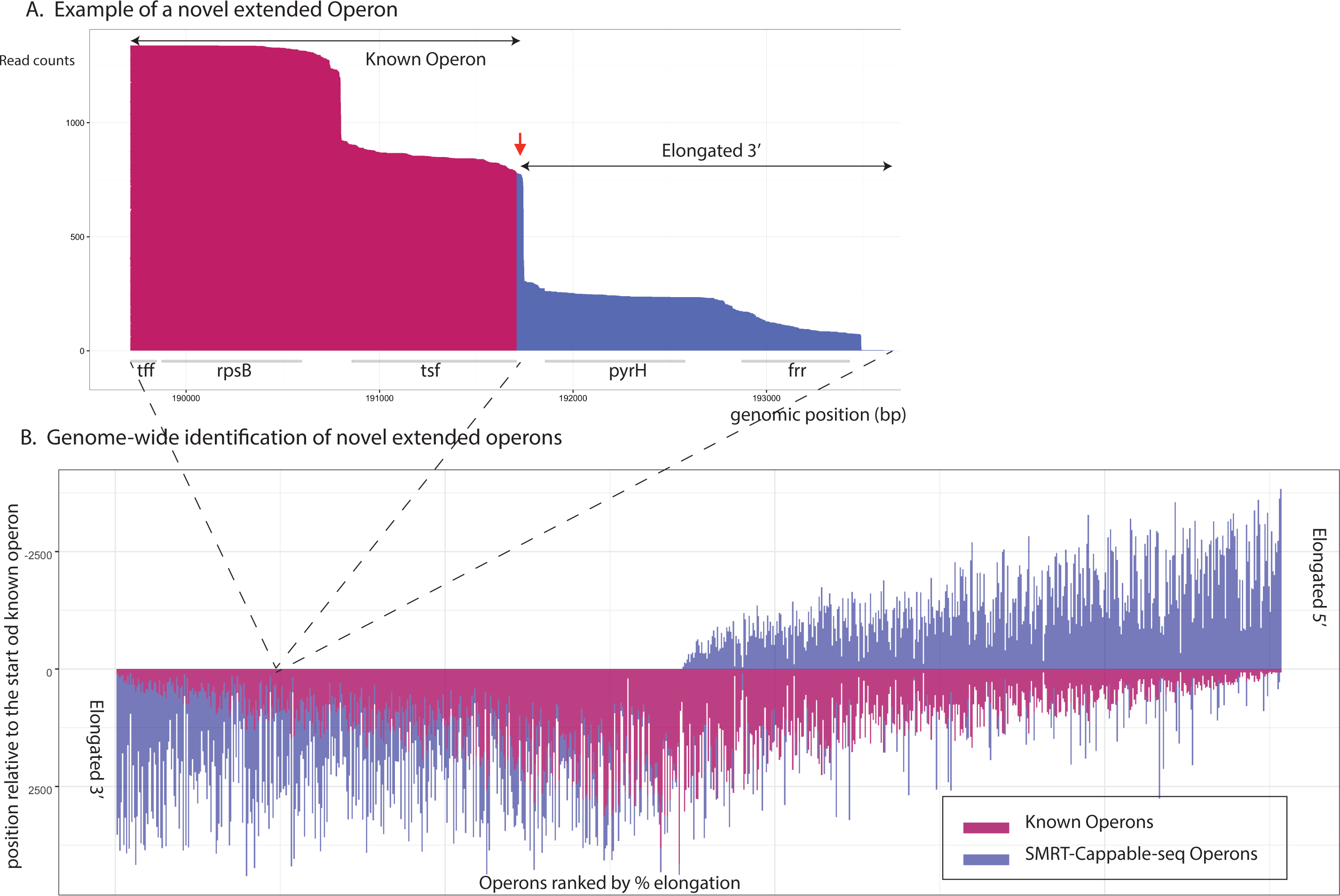
Extended Operons in *E. coli.* **A**. An example of the previously described tff-rpsB-tsf operon elongated at its 3’ end with two additional protein coding genes: pyrH and frr. The X-axis represents the position (in bp) on the reference genome (NC_000913.3) and the Y axis represents individual mapped reads ordered by read size in ascending order. For clarity only reads from the TSS (position at 189712) are shown. The red and blue parts of the reads correspond to the previously annotated operon and the extended operon respectively. Red arrow indicates the previously described Rho-independent terminator. tff encodes a putative small RNA. The product of pyrH is involved in nucleotide biosynthesis, while the products of rpsB, tsf and frr are involved in translation. **B**. Genome-wide distribution of elongated operons with additional gene(s). Elongated operons are defined by at least one SMRT-Cappable-seq read that covers the entire known operon and extends it to include at least one additional fully covered gene. Only the longest known operons were used, and sub-operons fully included in the longest operons were excluded from the analysis. Each vertical line corresponds to the positions of the extended operons (in bp) with the portion of the previously annotated operons in pink and the extended operons in blue. Positions are relative to the 5’end of the annotated operon with 0 being the TSS of the annotated operons. In total there are 883 RegulonDB operons extended by SMRT-Cappable-seq from either 5’ or 3’ end.

Summing up, our results demonstrate the ability of SMRT-Cappable-seq to comprehensively identify full-length primary transcripts at base resolution. By essentially targeting primary transcripts, SMRT-Cappable-seq is ideally suited to analyse transcriptional output and operon structures in bacteria.

### Extension of annotated operons

To take full advantage of the unique abilities of SMRT-Cappable-seq to obtain long-range continuity of transcripts, we simultaneously analyzed the positions of 5’ and 3’ ends of reads representing unique transcripts in *E. coli.* Surprisingly, we observed that a large fraction of reads are notably longer than the annotated operons in RegulonDB. These long reads often include additional genes that are indicative of extended operons.

To systematically evaluate the fraction of extended operons containing additional genes, we compared transcripts defined from the combined SMRT-Cappable-seq libraries with known operons from RegulonDB [18]. If at least one SMRT-Cappable-seq read fully covers and extends a RegulonDB operon by one or multiple gene(s) on the 3’, 5’ or both ends, this RegulonDB operon is defined as extended (**Material and Methods**).

Using this definition, we found that 34% (883 operons) of annotated operons are extended (**Table S4 and Figure 2B**)). This percentage represents a remarkable large number of new operons considering decades of studies done in *E. coli.*

For example, the well-described tff-rpsB-tsf operon was thought to only contain the tff attenuator and two genes coding for essential components of the translational machinery, the S2 protein (rpsB) and the translation elongation factor EF-TS (tsf) [19]. Furthermore, a Rho-independent TTS has been defined at the 3’end of the operon downstream of the tsf gene (**Figure 2A**) [19]. SMRT-Cappable-seq confirms the existence of both the known tff-rpsB-tsf operon and the TTS. Additionally, SMRT-Cappable-seq uncovers transcripts containing two additional genes, the pyrH and frr genes encoding for the UMP kinase and the ribosome recycling factor respectively (**Figure 2A**). This result demonstrates the existence of an additional longer operon encompassing the rpsB, tsf, pyrH and frr genes. Synteny analysis of the region across bacterial species has revealed that the gene arrangement of the region spanning rpsB, tsf, pyrH, and frr is entirely conserved between *P. aeruginosa* and *B. subtilis [20]*, consistent with the existence of a functional and conserved operon containing all 4 protein coding genes. Finally, the inclusion of the pyrH gene into the operon encoding three proteins involved in translational processes provides an experimental evidence indicating that the UMP kinase pyrH, is also involved in translational processes.

With new polycistronic operons containing additional genes, SMRT-Cappable-seq represents a powerful technique to identify full length transcripts and transcriptional boundaries. Because genes implicated in same metabolic pathways are often found on the same operon [1], SMRT-Cappable-seq also provides a powerful strategy to predict with high confidence, metabolic pathways and gene function in bacteria. While E. coli is one of the best annotated prokaryotic species, it also has a total of 924 genes of yet unknown or predicted function (based on RegulonDB annotation) [18]. From our dataset, 110 of these unknown genes are now found together with genes with known function in the extended operons. For example, two previously unknown genes yedJ and yedR are found in the same operon with dcm and vsr which encode DNA-cytosine methyltransferase and DNA mismatch endonuclease, respectively. Accordingly, we predict the yedJ and yedR genes to be involved in processing DNA modification or DNA repair.

### Novel operon and transcriptional context

Extended operons correspond to only a subset of differentially annotated operon in SMRT-Cappable-seq datasets. With the ability to link the 5’ to the 3’ end of transcripts, SMRT-Cappable-seq unravels a number of sub-operon within operons leading to transcripts with novel combination of genes.

By analogy with transcripts isoforms as a result of splicing in eukaryotes, we describe such modular usage of operons as variants of operon. To avoid an overestimation of the number of operons, we use a conservative definition of operons for which a define TSS and a define TTS can be found and if no TTS can be found, the longest read is used to define operons (**Material and Methods**).

We found 2347 operons are encoding unique combination of genes and amongst those, 840 are novel compared to RegulonDB **(Table S3 and Supplementary text)**. SMRT-Cappable-seq operons have an average of 2.1 genes per operon compared to 1.4 in RegulonDB suggesting that polycistronic operons are more prevalent than previously thought: around 30% of the defined operons start within another operon (see **Figure S4** for examples).

To assess the extent of operon variants in *E. coli*, we define Transcriptional Context (TC) as the unique combination of genes within an operon, and investigated for each annotated gene, the number of unique TC the gene is found in. We found that around 50% of genes analyzed are in more than one TC with some genes being found in, as many as 15 TCs (**Figure 3A and C**). Genome-wide, SMRT-Cappable-seq identifies 2370 more genes in multiple TCs than those annotated in RegulonDB. The Ribosome-recycling factor frr gene exemplifies the extent of TC modularity with frr being found to be transcribed in eight different TCs (**Figure 3B, Table S5**) each of them having a different gene configuration, whereas RegulonDB and previous frr gene literature describes only the transcript containing the frr gene alone [21].

**Figure 3:**
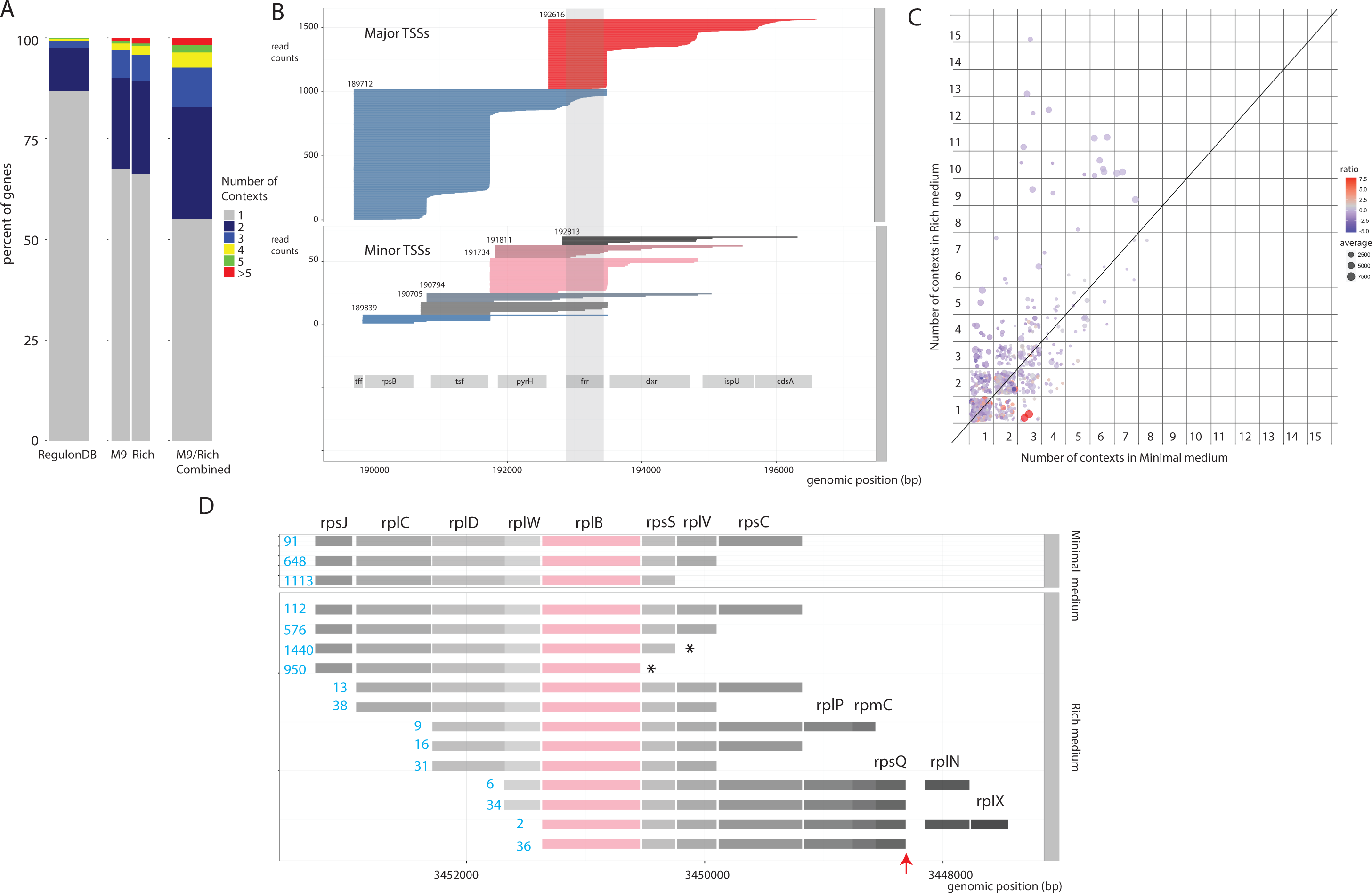
Transcriptional context. **A**. Distribution of genes according to the number of transcriptional contexts from RegulonDB (left barplot), SMRT-Cappable-seq (Center barplots) and combined SMRT-Cappable-seq dataset (Right barplot). Only genes having at least one context are considered. The number of transcriptional contexts for a given gene X is the number of unique combinations of genes cotranscribed with gene X. **B**. Individual reads (Rich condition) mapping across the frr gene ordered by TSS location and decreased read size. Read colors represents TSS positions. Reads derived from major and minor TSS are divided into two panels offering different scale for clarity. The frr gene is transcribed from 8 different gene contexts when the cells are grown in Rich medium. These contexts are altering the gene content of the transcripts encoding the frr protein. The 8 possible combinations are: rpsB-tsf-pyrH-frr, tsf-pyrH-frr-dxr, tsf-pyrH-frr, pyrH-frr-dxr, pyrH-frr, frr-dxr-ispU-cdsA, frr-dxr-ispU and frr alone. RegulonDB only annotates frr alone. **C**. Number of contexts in M9 medium (x axis) compared to the number of contexts in Rich medium for each selected genes. The point size represents the average expression level between the two growth conditions while the color represent the ratio between the expression level in Rich medium compared to M9 medium. **D**. Example of the rplB gene coding for the 50S ribosomal subunit protein L2 showing a change in the number of transcriptional context from 13 contexts (Rich medium) to only 3 contexts (M9 medium). * indicates the SMRT-Cappable-seq defined TTSs downstream of rpsS and rplB, and red arrow indicates the previously known TTS downstream of rpsQ. The number of reads fully covering the operon is shown in blue.

Interestingly, TC can be condition dependent: of genes expressed during growth in both M9 and Rich media, 35% display differences in TC. Those genes tend to be found in a greater variety of TC in Rich compared to M9 medium (**Figure 3C**). For example, the rplB gene coding for the 50S ribosomal subunit protein L2 is transcribed in 13 different TCs when the cells are grown in Rich medium compared to only 3 in minimal medium (**Figure 3D**). Genes with more TCs in Rich medium (**Table S5**) are significantly enriched in biological processes involved in translation (GO analysis, 12-fold enrichment, p.value = 1.62E-09 with Bonferroni correction for multiple testing). These results demonstrate that genes within multiple TCs are more common in bacteria than previously thought highlighting a novel level of gene control that has not yet been thoroughly assessed yet.

### Read-through at transcription termination site (TTS)

Next, we investigated the possible mechanisms underlying the existence of the large number of operon variants found by SMRT-Cappable-seq. We found that many of the sub-operons and extended operons that we identify are the consequence of extensive read-through at TTS. The concept of read-through is not new and has been demonstrated for some operons [22-24], synthetic constructs [25] and described for convergent operons [26]. Nevertheless, the extent of readthrough across prokaryotic transcriptome and the link between TTS readthrough and operon variants has never been explored.

SMRT-Cappable-seq highlights the pervasiveness of the *in-vivo* readthrough of the entire transcriptome with 75% of TTS having at least 5% of read-through transcripts (**Supplementary text**). TTSs overlapping with experimentally validated TTSs show essentially similar behavior with 70% of known terminators having read-through (**Figure 4B**). The degree of read-through largely depends on the TTS, in some cases as high as 50% of the transcripts extending across sites (**Figure 4B and Figure 5A**). Interestingly, the extension in the majority of the cases is longer than 100 bp on average (**Figure 4C**) and around 40% of the read-through transcripts include additional gene(s), which indicates that the downstream extension is functional. As a result, display of transcripts sharing the same TSS collectively forms a staircase pattern originating at the 3’ end (**Figure S5**).

**Figure 4:**
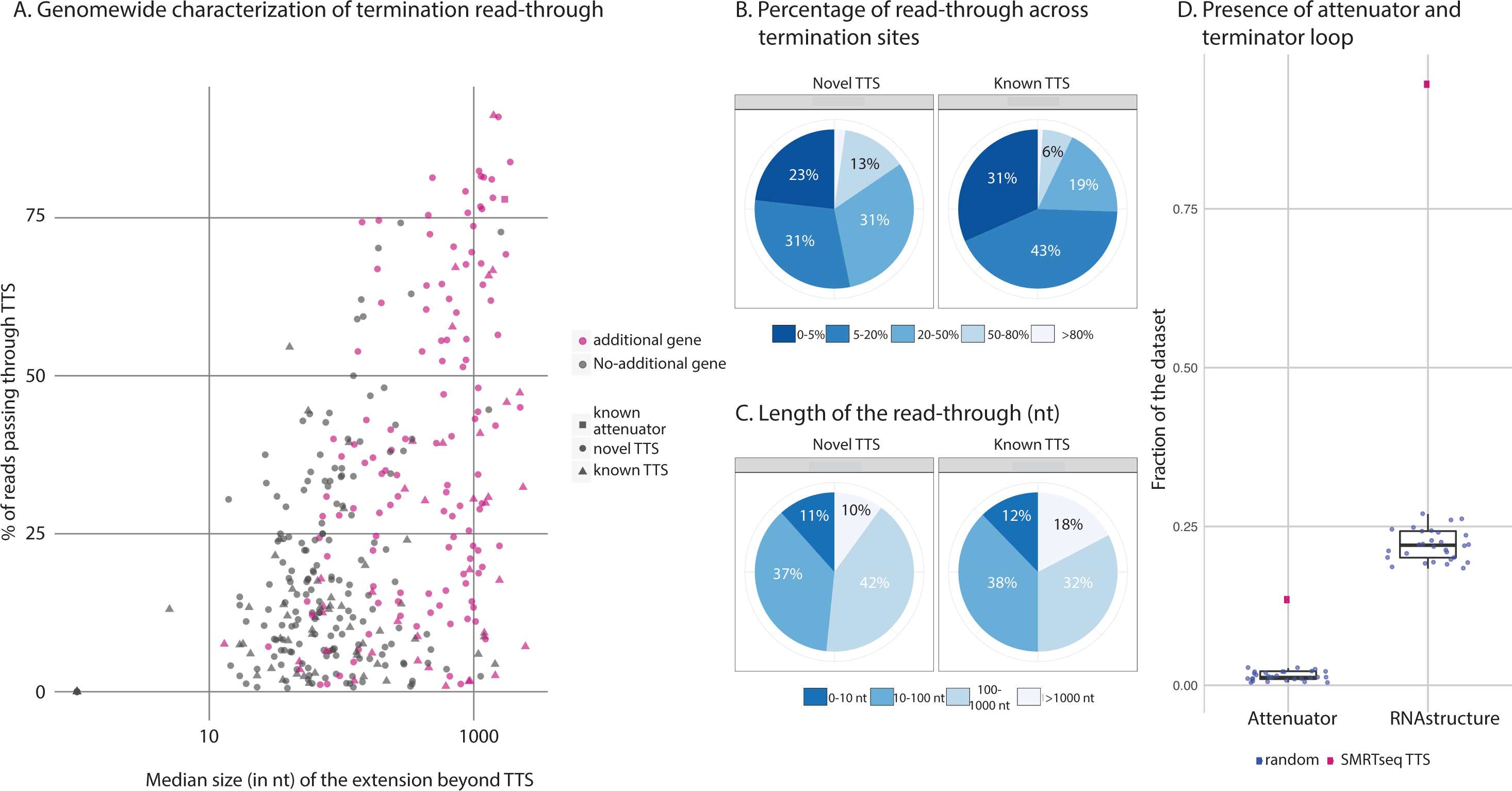
The vast majority of termination sites have read-through. **A**. Relation between the percentage of read-though (Y axis) and median size of the region (in bp) located 3’ of the TTS (X axis) for every transcripts with a defined TSS in M9 medium. TSSs with read-through containing additional gene(s) are in pink. **B**. Pie chart showing the range of percentage of read-through for novel (left panel) and known (right panel) terminator sites in M9 medium. A TSS-TTS pair is identified as operons with a defined TSS and TTS. In total, 347 unique TTS were defined by SMRT-Cappable-seq from a total of 408 TSS-TTS pairs. Here read-through means that the 3’ end of the read exceeds at least 50 bp of the termination site. In 305 out of 408 TSS-TTS pairs (Table S2), the read-through is above 5%. For 67 of the 98 previously known TTS (red), the read-through is above 5%. **C**. Pie chart showing the range of median size in bp of the 3’ extended region passing the termination site in M9 medium. 151 out of 408 termination sites contain additional genes. **D**. Percentage of termination sites in minimal medium (red) and random positions (blue) with predicted attenuator structures (left panel) and predicted termination loop (Pasific algorithm [32]) (right panel). 55 out of 408 termination sites have a predicted attenuator structure, and 386 of 408 termination sites have a predicted terminator structure.

Comparison between growth conditions reveals that, in some cases, the degree of read-through is conditionally dependent (**Figure 5A and B, Figure S6A**), consistent with TTS serving as a major modulator of gene regulation. For example, the prs-dauA operon contains a TTS between the prs and dauA gene that shows 91% read-through in Rich medium growth as opposed to only 43% in M9 condition (**Figure 5A**). This change in the degree of read-through results in the apparent up-regulation of the dauA gene in Rich condition despite similar expression levels at the TSS. The dauA gene encodes for the C4 dicarboxylic acid transporter, which mediates succinate transportation. Previous studies have shown that the acetate accumulation rate is faster in Rich compared to M9 medium [27], which leads to an acidic pH. As a result, succinate has to be excreted from the cell, which requires the expression of the dauA gene [28].

**Figure 5:**
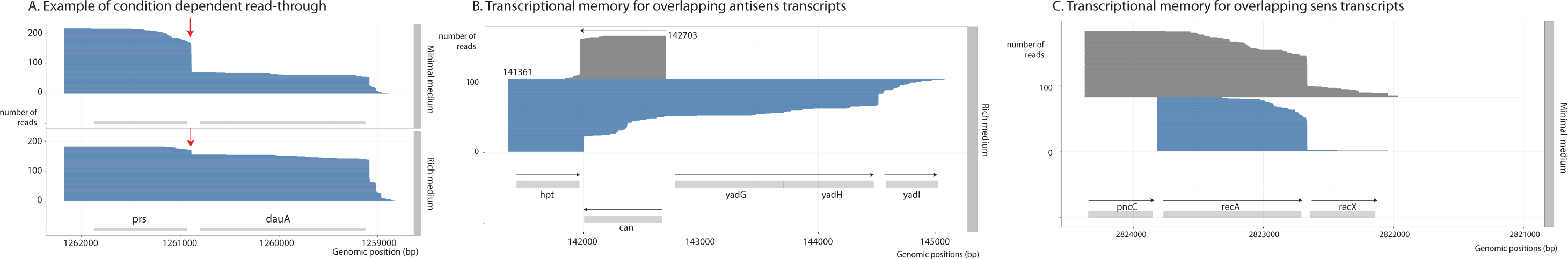
Transcriptional coordination. **A**. And example of the prs-dauA operon containing a previously known TTS (red arrow), where the rate of read-through is condition dependent. The TTS between the prs and dauA gene shows 91% read-through in Rich medium as opposed to 43% read-through in M9 medium. As a result the dauA gene is up-regulated in Rich medium compared to M9 growth condition despite comparable expression level between conditions for its co-transcribe prs gene. **B**. An example of a transcriptional coordination where for the same termination site, the majority (90%) of antisense transcript coding for the can gene terminates while the majority (78%) of the transcripts coding for hpt has read-through. **C**. Overlapping transcripts in the same orientations. 26% read-through vs 5% read-through.

Therefore, the control of read-through of the TTS is a possible mechanism to enable the bacteria to adjust their enzyme levels quickly to the fast changing environment.

Furthermore, we found that the degree of read-through of a given terminator can vary according to the TSS, a phenomena that we term “transcriptional coordination” because the transcription machinery “coordinates” the location of the transcription start site with the TTS read-through. We found cases of transcriptional coordination in overlapping antisense transcripts suggesting that both transcripts are using the same forward/reverse termination site but the extent of read-through is different. This difference in readthrough is expected given that antisense transcripts have different sequence composition (**Figure 5B**). More interestingly, we also identified a few cases of transcriptional coordination in overlapping transcripts on the same strand (Figure 5C). Accordingly, we hypothesize that the overall structure difference between transcripts initiated at different TSS maybe a mechanism for such coordination.

Next we examined whether this pervasive read-through across TTS is the consequence of “weak” terminators. For this, we predicted the termination loop structure, and consistent with the previous work, we found no correlation between the free energy of the RNA secondary structure and the degree of TTS read-through (**Figure S2C**) [29, 30]. Furthermore, a similar motif profile containing the classic loop structure was observed for TTS datasets with both low and high read-through (**Figure S2B**). The polyU characteristic of the region of the termination sites is present for both high and low read-through TTS, however, and consistent with previous findings, we found that the polyU tract is better defined in the low read-through TTS dataset (**Figure S2B**) [31]. Collectively, these results indicate that the structure of the termination loop tends to be well-conserved with a better defined polyU track for the stronger TTS.

Next, we examined whether regions flanking the loop can be another determinant of the read-through level. Previous studies have described riboswitches leading to two mutually exclusive RNA structures, one of which forms a transcriptional terminator and results in transcription termination, and the other forms an antiterminator that allows read-through across the terminator [3, 17]. While riboswitches have been extensively studied in the past, they have primarily been identified in the 5’ UTRs of operons to modulate the commitment of the RNA polymerase to either fully transcribe or prematurely abort transcripts [17]. We analysed the TTS found by SMRT-Cappable-seq using a published algorithm for riboswitches prediction and found that 13.5 % of the TTS are preceded by predicted attenuator structures that resemble riboswitches [32] (**Figure 4D**). That is a 10 fold enrichment of attenuator structures at SMRT-Cappable-seq TTS compared to randomly selected sequences suggesting that the read-through is modulated via such structures.

As most of the SMRT-Cappable-seq TTSs are located 3’ of a transcribed gene, we hypothesized that riboswitch-like structures are also modulating read-through in the 3’ UTRs of transcripts after the polymerase has transcribed one or several gene(s). Under this hypothesis, riboswitch-like structures are not only controlling the commitment of the RNA polymerase to transcribe operons but are also involved in a more fine-tune control of the gene compositions of operons.

To test this hypothesis, we studied the known dapA-bamC Rho-independent terminator that shows 44% and 14% differential read-through in M9 and Rich medium respectively (**Figure S6A**), a differential read-through that we confirmed by qPCR. A Riboswitch structure was predicted 5’ of the TTS [32] (**Figure S6B**). We deleted the sequence predicted to be riboswitch-structure in the bacterial genome [33] and measured the level of read-through in both growth conditions using qPCR. We found that deleting the riboregulator sequence, while it does not affect the read-through in M9 growth condition, it significantly (p-value<0.005) increased the read-through in Rich medium (**Figure S6C and D**). This suggests that this riboswitch-structure controls the conditional-regulated termination.

Taken together, these results demonstrate that a large number of operon variants are resulting from read-through at TTS. With the majority of operon termination sites having substantial read-through, this study redefines termination sites as a major point of control of gene expression. Here we show that the termination sites of operons have similar modular capacities to create polycistronic transcripts with variable gene compositions. Our results provide experimental evidences that a large fraction of bacterial gene expression is regulated by conditional-regulated termination, possibly via riboswitches structures [32].

## Discussion

By highlighting for the first time a comprehensive view of the full-length transcriptional landscape, this study provides an important resource for functional annotation and the understanding of gene regulation in bacteria and microbiomes. Applications in microbiomes are particularly appealing since SMRT-Cappable-seq will be able to uncover partial or complete metabolic pathway by phasing functionally related genes on the same sequencing reads.

This unique ability to phase transcripts over long distances has also revealed the complexity of operon structures, complexity driven by both the regulation of transcription initiation and the modularity of transcription termination notably termination read-through. While a number of studies have accumulated read-through evidences for synthetic constructs or isolated transcripts [22, 25], this study demonstrates the pervasive and modular nature of read-through of termination sites across bacterial transcriptome. This read-through represents a powerful mechanism to control elongation of operons and include additional genes. In some cases, this control can be condition dependent and lead to the net up or down regulation of gene expression despite constitutive promoter activity.

Such modularity in the degree of transcription termination leads to a spectrum of sub-operons within larger operons and the ability of bacteria to transcribe genes in an array of transcriptional contexts. In the case of RNA-seq these transcriptional contexts are hidden and their importance have been underestimated. Yet, several studies have demonstrated that not only the level of gene expression is important for function but also the larger transcriptional context by which a gene is expressed. For example, a recent paper has shown that co-translation of genes on the same transcripts influences the proper arrangement and folding of complex protein structures [34] highlighting the importance of relating gene expression in the context of other genes on the same transcripts.

---

**Supplementary Figure 1:**
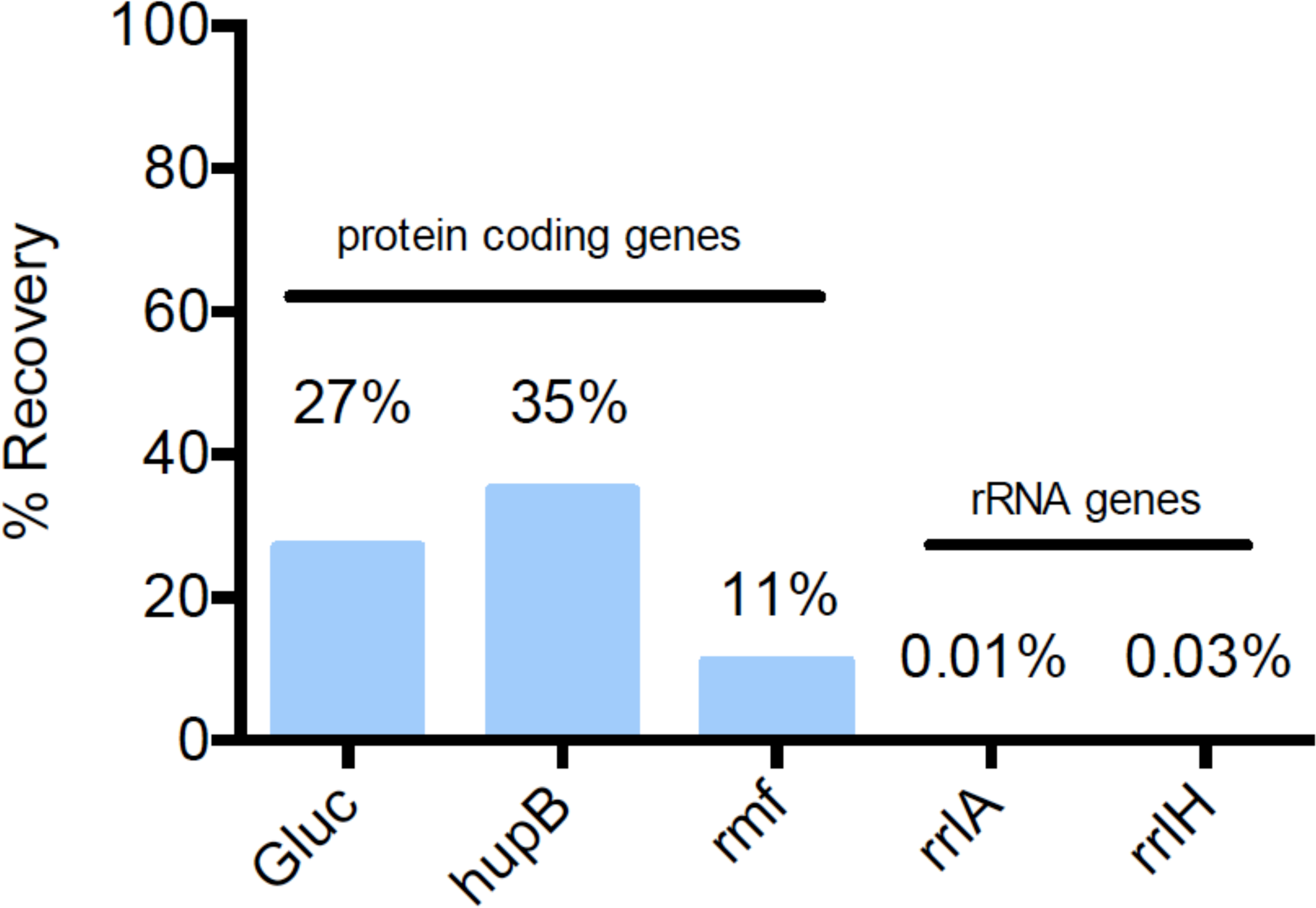
Recovery of mRNA and rRNA by SMRT-Cappable-seq. The mRNA levels of protein coding genes (Gluc, hupB and rmf) and rRNA genes (rrlA and rrlH) were measured by qPCR using the cDNA obtained from *E. coli* grown in M9 medium. The recovery rates (in %, Y axis) was calculated as the amount of mRNA in the enriched fraction (after streptavidin) divided by the amount of mRNA in the control fraction (no streptavidin enrichment). Gluc is an *in-vitro* transcribed mRNA spiked in the *E. coli* total RNA as positive triphosphorylated control. HupB and rmf are endogenous protein coding genes representative of primary transcripts. rrlA and rrlH are rRNA representative of processed transcripts.

**Supplementary Figure 2:**
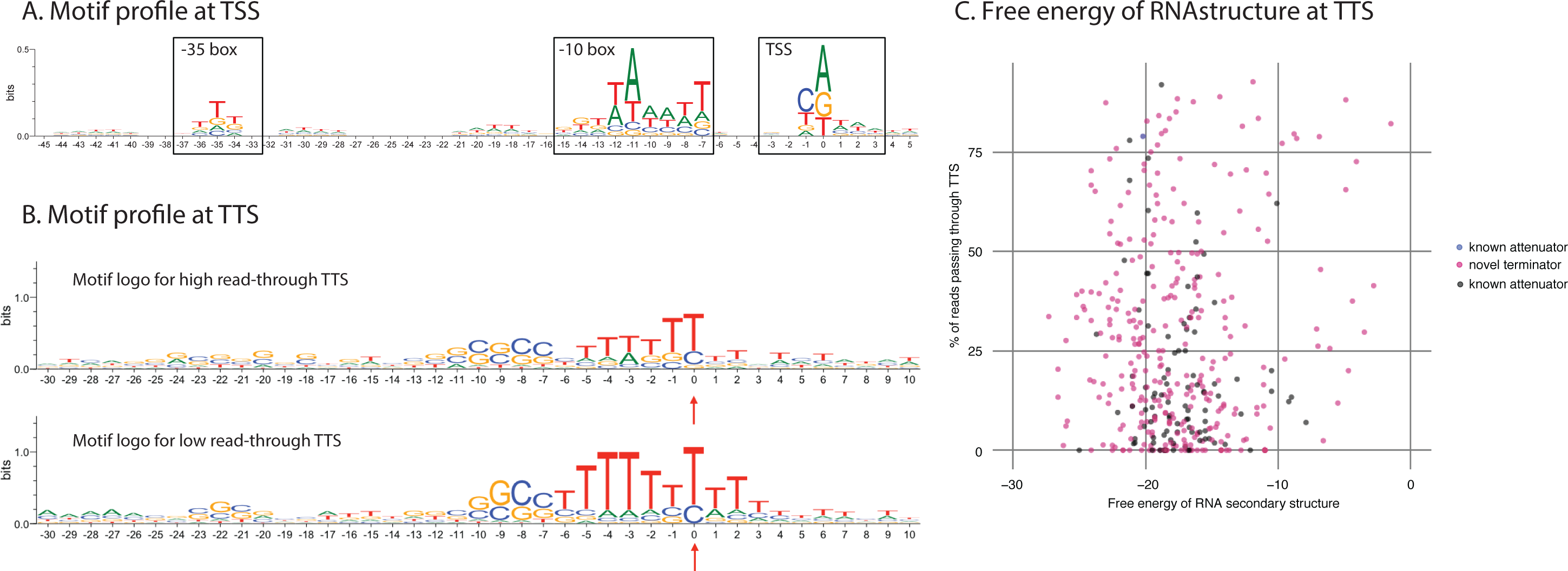
Motif and RNA structure analysis for TSS and TTS in M9 medium. **A**. Motif logos for SMRT-Cappable-seq defined TSSs. X-axis: Position 0 corresponds to TSS, negative and positive values correspond to positions upstream and downstream of TSS respectively. Y-axis : information content expressed in bits. **B**. Motif logos for TTSs (top) that have high (more than 25%) read-through and for TTSs (bottom) that have low (less than 25%) read-through. Red arrow indicates the termination site. X-axis: Predicted TTS is located at position 0, negative and positive values correspond to positions upstream and downstream of TTS respectively. Y-axis: information content expressed in bits. **C**. No correlation (corr = 0.035) between the degree of read-through of the TTS and the free energy of the RNA structure (x axis) of the TTS can be observed.

**Supplementary Figure 3:**
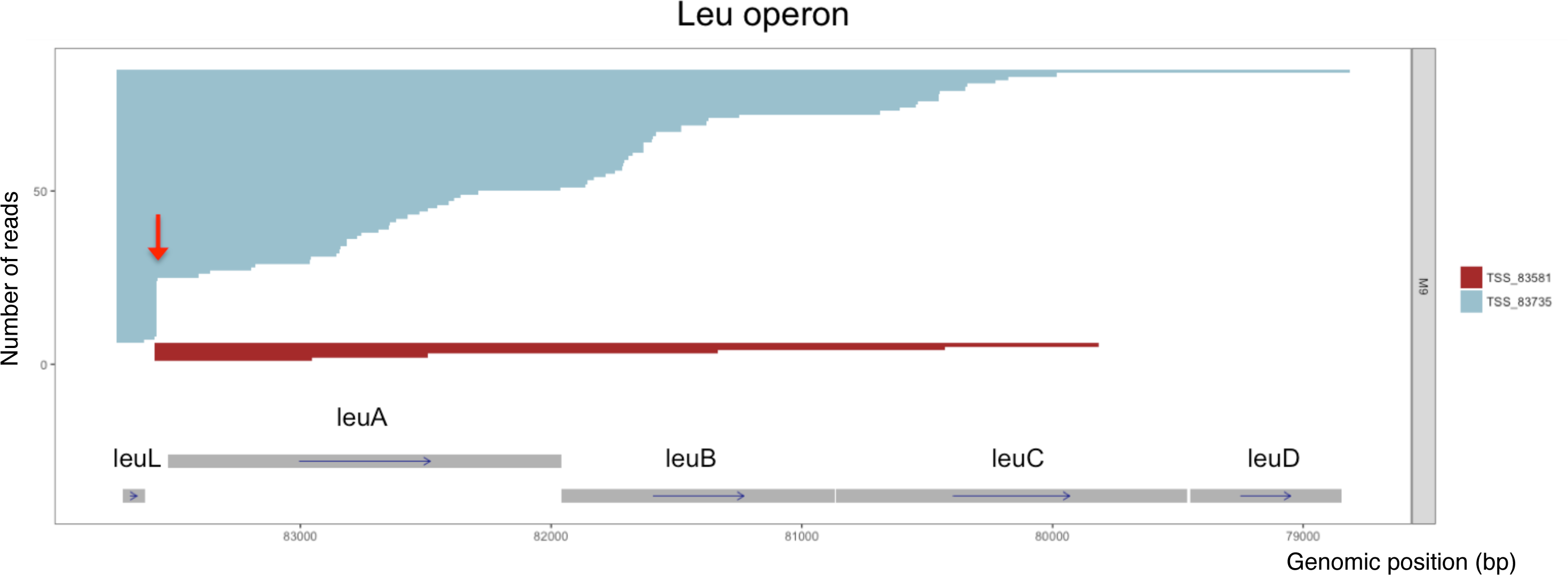
Structure of the Leu Operon. Individual mapped reads ordered by TSS location and read size in the Leu operon locus. The red arrow denotes the termination site controlled by a riboswitch. Reads in blue represent transcripts originated from the TSS upstream of the leader peptide leuL (pos = 83735). Reads in red represent transcripts originated from the TSS downstream of the leader peptide (pos = 83581).

**Supplementary Figure 4:**
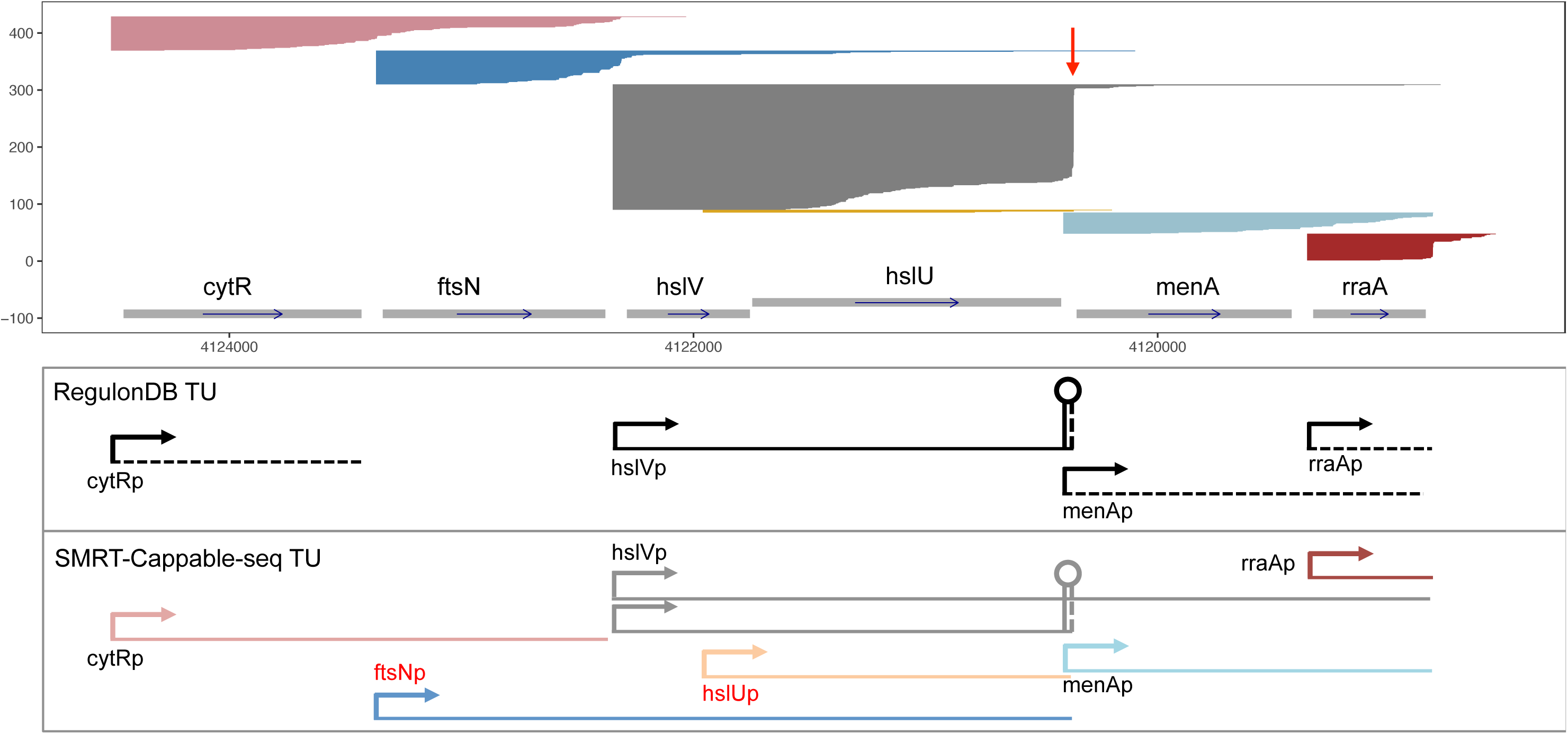
SMRT-Cappable-seq read profile in Rich condition. Schematic representation of previously annotated operons from RegulonDB database (dash line indicates weak evidence, and solid line indicates strong evidence) and operons defined by SMRT-Cappable-seq. There are four annotated operons covering the coding genes in the shown genome region, three of which are also defined by SMRT-Cappable-seq (hslV-hslU, menA-rra, rra). SMRT-Cappable-seq additionally identified four extended operons. Two of them (ftsN-hslV-hslU and hslU) have previously unidentified promoters and 5’ genes. The other cytR-ftsN has the same promoter as the previous cytR unit, but includes an additional gene at the 3’end. Another extended operon is composed of the hslV-hslU-menA-rraA genes due to the read-through at the hslV-hslU terminator. Red arrow indicates the previously known TTS for the operon.

**Supplementary Figure 5:**
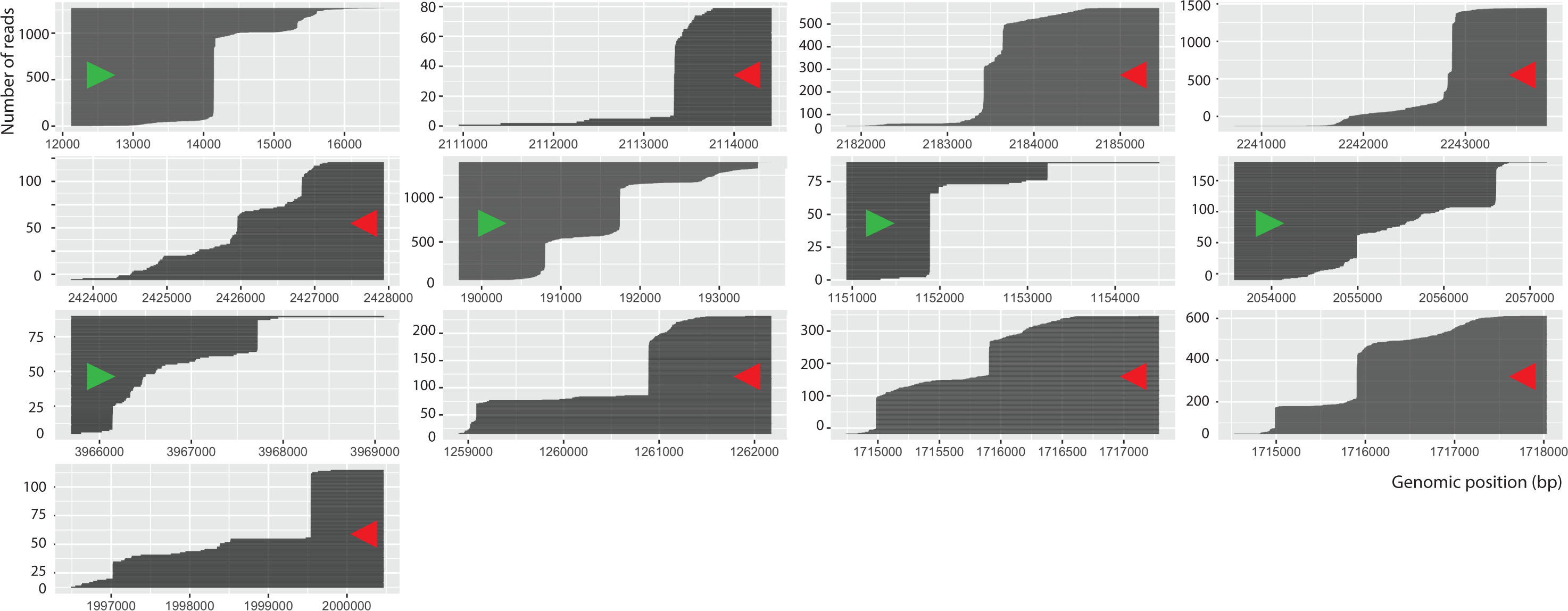
Read-through across TTS leads to operon variants. Examples of operon variants generated from the same TSS as a result of read-through of termination sites. Data shown is from M9 growth condition. Arrows denote the direction of transcription on the forward strand (green arrow) and reverse strand (red arrow).

**Supplementary Figure 6:**
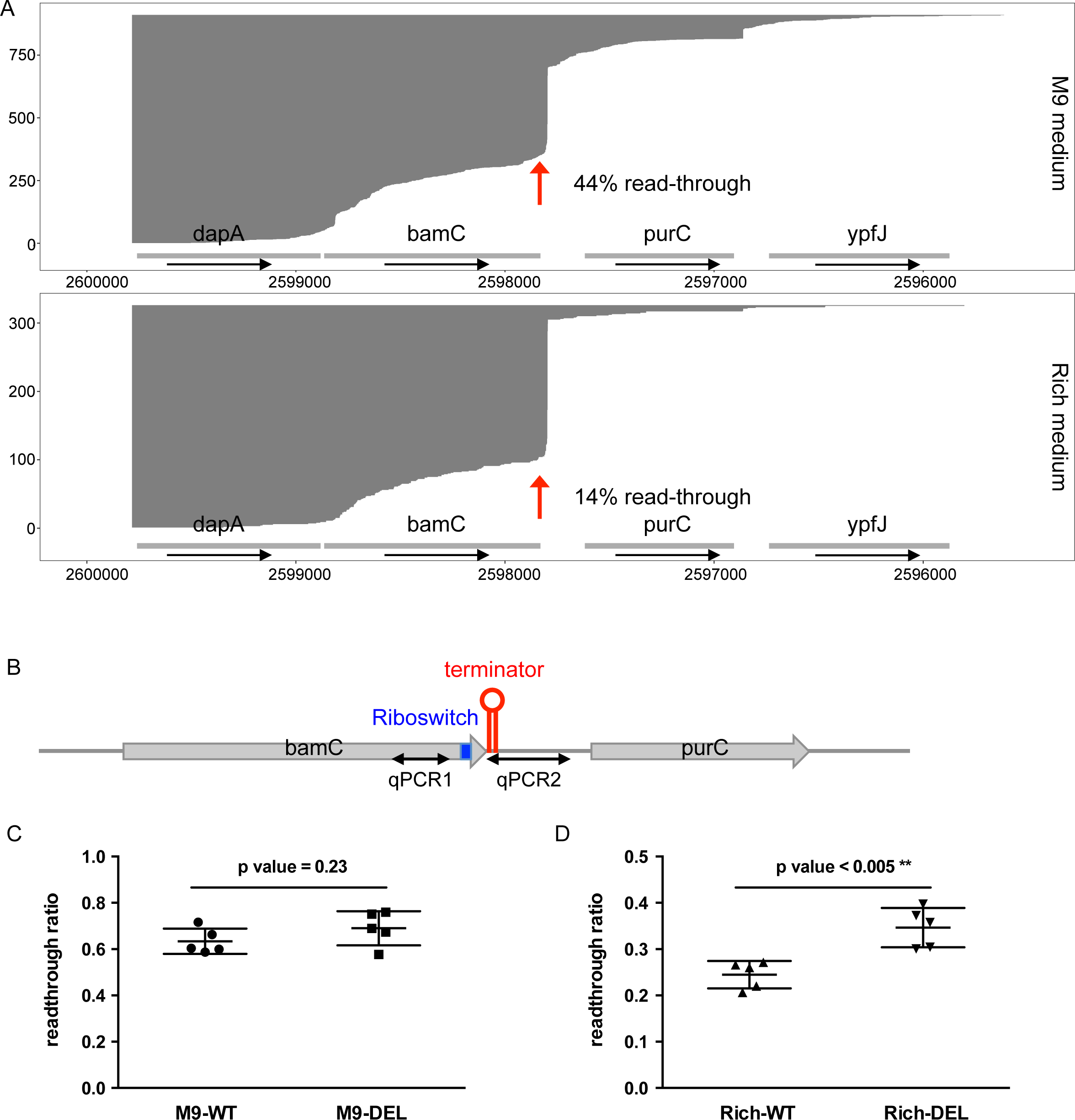
The riboswitch structure controls the condition-dependent read-through across the dapA-bamC termination site. **A**. An example of the dapA-ypfJ operon containing the previously known dapA-bamC TTS (red arrow), where the rate of read-through is condition dependent. **B**. Schema of this TTS location. The Riboswitch structure (blue) that locates 10 bp upstream of the termination site (red) is deleted to examine the role of this regulatory region in the control of transcription termination. **C. D**. The level of read-through across the TTS of wild-type (WT) and deletion strain (DEL) were measured for bacteria grown in both M9 and Rich medium by qPCR. The qPCR1 primers amplify a upstream region of the predicted riboswitch site. For qPCR product 2, the forward primer binds to the 5’ end of the known dapA-bamC TTS while the reverse primers binds to the downstream region of the TTS. Therefore, read-through ratio was calculated as the amount of qPCR2 product divided by qPCR1 product. Data shown are the means ± SD from five independent repeats. Unpaired t-test was used to determine significance. The read-through in DEL is significantly higher than in WT in Rich condition.

**Supplementary Figure 7:**
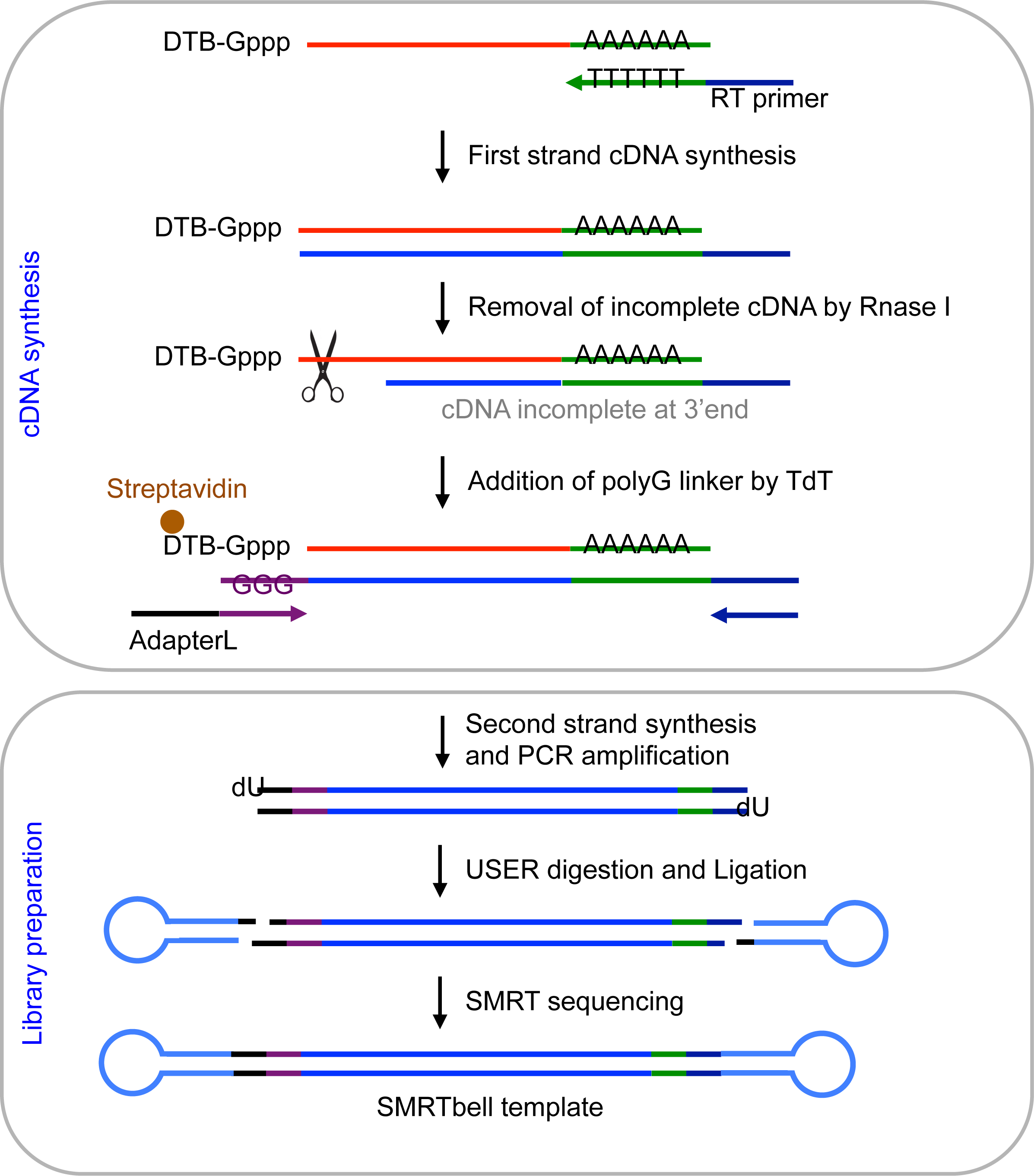
Schema of the SMRT-Cappable-seq library preparation. First strand cDNA is synthesized using RT primer containing dT sequence. The RT step is followed by Rnase I treatment to remove incomplete cDNA fragment. Terminal transferase is then used to add a polyG tail at the 3’ end of the cDNA. Second strand cDNA is made and further amplified by PCR. The dUracil in the primer is removed using USER to create sticky end, and PacBio adapters are ligated to the amplified fragments to generate SMRTbell template. SMRT sequencing of the resulting SMRTbell templates provide the genome-wide definition of full-length transcripts at base resolution.

## Supplementary Tables

**Supplementary table 1: Sequences of primers used in this study.**

**Supplementary table 2: TTSs defined by SMRT-Cappable-seq under M9 minimal condition**

See Supplementary text for details.

Column 1: Genome version

Column 2: start, which shows TSS for + strand OR 3’ boundary of the clustered TTS for - strand

Column 3: end, which shows TSS for - strand OR 3’ boundary of the clustered TTS for + strand

Column 4: 5’ boundary of the clustered TTS (equals to Column 3 if there is only one TTS in the cluster)

Column 5:.

Column 6: strand

Column 7: The number of reads starting at the TSS position

Column 8: The number of reads starting at the TSS and ending in the TTS cluster

Column 9: The number of reads having read-throughs across this TTS

Column 10: Median size of the 3’ extended region (bp).

Column 11: TTS types (novel TTS, known terminator or known attenuator).

Column 12: Name of the nearest known terminator.

Column 13: The position of the known terminator (center).

**Supplementary table 3: Operon Annotation**

Operons defined by SMRT-Cappable-seq under both M9 and Rich condition were compared with the previous annotation from RegulonDB.

Column 1: List of genes (separated by |) in the SMRT-Cappable-seq defined operon

Column 2: Operon strand

Column 3: Comparison with RegulonDB. See Supplementary text for details.

Column 4: List (Separated by semi-colon) of previously annotated RegulonDB operons that overlap with the SMRT-Cappable-seq operon

**Supplementary table 4: Extended Operons in Regulon DB**

See Supplementary text for details.

Column 1: List of gene (separated by |) in RegulonDB annotated operons

Column 2: List of genes contained by SMRT-Cappable-seq transcripts

**Supplementary table 5: Transcriptional Context (TC)**

Column 1: Gene ID

Column 2: The number of TC that the gene is found in under M9 condition

Column 3: The number of TC that the gene is found in under Rich condition

## Acknowledgements

The authors would like to thank Rich Roberts, I. Schildkraut, L. Raleigh, M. Berkmen, B. Jack, T. Carlow, L. McReynolds, H. Runz and D. Wang for critical comments. Disclosure declaration: L.E. and B.Y. are employees of New England Biolabs. M.B and T.C. are employees of Pacific Biosciences of California.

## Author contributions

Y. developed the methodology and performed the experiments. M.B. and T.C. assisted with the library preparation and performed the SMRT sequencing. B.Y. and L.E. analyzed the data. L.E. and B.Y. wrote the manuscript. All authors approved the final manuscript.

## Declarations

Raw Sequencing reads have been deposited at the European Nucleotide Archive. The computer scripts used for the data analysis in this study are freely available on github.

## Abbreviation

TSS: transcription start site
TTS: transcription termination site
TC: transcriptional Context
TU: transcription unit
SMRT: Single Molecule, Real-Time
RT: reverse transcription
rRNA: ribosomal RNA
M9: minimal growth medium

## Material and Methods

### Data availability

Sequencing data have been deposited at the European Nucleotide Archive. For review the data could be accessed through: ftp://ftp.neb.com/unpub/yan/

### Code availability

The computer scripts used for the data analysis in this study are freely available on github (https://github.com/elitaone/SMRT-cappable-seq).

### Growth conditions and RNA isolation for SMRT Sequencing

*E. coli* k-12 strain MG1655 was grown at 37°C in M9 minimal medium with 0.2% glucose and Rich medium (per liter 10 g Tryptone, 5 g Yeast Extract, 5 g sodium chloride, pH 7.2), respectively. The culture was grown to late log phase (0D600 between 0.55-0.6). 2 volumes of RNAlater (Life Technologies) was added to the culture and saved at 4°C overnight. The RNA was extracted using RNAeasy Midi kit (Qiagen). The isolated RNA had a RIN score above 9.0 as determined by Bioanalyzer (Agilent), and was used for cDNA synthesis, library preparation and SMRT Sequencing.

### Enrichment of full-length prokaryotic primary transcripts (**Figure 1A**)

The capping reaction was performed on total RNA as described in [4] with the following modifications: 5 μg of *E. coli* RNA was incubated in the present of 0.5mM DTB-GTP (New England Biolabs) and 100 units of Vaccinia Capping Enzyme (New England Biolabs) for 0.5h at 37°C in 100 μl reaction volume. In order to measure the recovery of triphosphate transcripts, 1ng of *in vitro* synthesized Gluc (Gaussia Luciferase) transcripts were mixed with the *E. coli* RNA for the capping and following reactions. The capped RNA was purified using AMPure beads.

Next, a polyA tail was added *in-vitro* by incubating the capped RNA in 50 μ! reaction volume with 20 units *E. coli* Poly(A) Polymerase (New England Biolabs) and 1mM ATP for 15min at 37°C. The capped and tailed RNA was purified using AMPure beads and eluted in 50 μl low TE buffer. 10 μl of the reaction (non-enriched RNA) was put aside and used as control.

The capped RNA was enriched using hydrophilic streptavidin magnetic beads (New England Biolabs). 40 μl of beads were prepared as described [4] and suspended in 40 μl Binding Buffer (see below for composition). 40 μl of capped and tailed RNA was incubated with the beads at room temperature for 30min on a rotator. The beads were then washed thoroughly three times with Binding Buffer and three times with Washing Buffer. To elute the RNA, the beads were resuspended in 26 μl Biotin Buffer and incubate at 37°C for 25min on a rotator, and 14 μl of Binding buffer was added and incubate for another 5min. The Biotin eluted enriched RNA was purified with AMPure beads and eluted in 20 μl Low TE.

Low TE buffer: 1mM Tris-HCl pH7.5; 0.1mM EDTA

Binding Buffer: 10mM Tris-HCl pH7.5; 2M NaCl; 1mM EDTA

Washing Buffer: 10mM Tris-HCl pH7.5; 250mM NaCl; 1mM EDTA

Biotin Buffer: 1mM Biotin; 10mM Tris-HCl pH7.5; 0.5M NaCl; 1mM EDTA

### cDNA synthesis for SMRT Sequencing (**Figure S7**)

20 μl of enriched RNA (SMRT-Cappable-seq) and 10 μl of non-enriched RNA (control)) were used in a 40 μl first strand cDNA synthesis reaction with 400 units of ProtoScript II Reverse Transcriptase (New England Biolabs) and 50 μM RT primer (**Supplementary Table 1**). Reactions were incubated at 42°C for 1h, then 50 units Rnase If (New England Biolabs) was added and incubated at 37°C for another 30min. The reactions were purified using AMPure beads and eluted in 20 μl Low TE.

The polyG linker was added to the 3'end of cDNA for second strand synthesis using Terminal transferase (TdT, New England Biolabs). The purified cDNA/RNA duplex samples were incubated with 10 units TdT and 2mM dGTP at 37°C for 30min. For the SMRT-Cappable-seq RNA, the reaction was purified using 30 μl hydrophilic streptavidin magnetic beads (New England Biolabs) as mentioned above but without Biotin buffer elution. The washed beads with the bound cDNA/RNA duplex were resuspended in 30 μl Low TE. For control RNA, the reaction products were purified using AMPure beads and eluted in 30 μl Low TE.

### SMRTbell library preparation and SMRT Sequencing (**Figure S7**)

Second strand cDNA synthesis was performed on both control and enriched first strand cDNA/RNA duplex samples with LongAmp HotStart Taq DNA polymerase (New England Biolabs) using Pac_oligodC20 and Pac_rev primers following the manufacturer’s instructions.

PCR cycle number optimization was used to determine the optimal amplification cycle number for the downstream large-scale PCR reactions as described in the standard iso-seq protocol (http://www.pacb.com/wp-content/uploads/2015/09/User-Bulletin-Guidelines-for-Preparing-cDNA-Libraries-for-Isoform-Sequencing-Iso-Seq.pdf). Second-strand cDNA was then bulk amplified with LongAmp HotStart Taq using Pac_for_dU and Pac_rev_dU primers, which contain an internal 5'-dUTP. Large-scale PCR products were purified with AMPure PB beads (Pacific Biosciences) and quality control was performed on a BioAnalyzer (Agilent). cDNA was then subjected to size fractionation using the Sage BluePippin system (Sage Science), collecting three size-bins: 1-2kb, 2-3kb and 3-6kb. Following size-selection, size-selected cDNA was re-amplified using LongAmp HotStart Taq. Purified, size-selected cDNA was eluted in 20ul Low TE and then digested using 1 unit USER (New England Biolabs) to create sticky ends. The the sticky ended cDNA for each size-bin was then ligated to μM hairpin adapters that contain complementary sticky ends to create SMRTbell templates using 2000 Units T4 DNA ligase (New England Biolabs). After incubation at room temperature for 1h for ligation, 100 Units Exonuclease III (New England Biolabs) and 10 Units Exonuclease VII (USB) were added into the ligation reaction to remove the incompletely formed SMRTbell templates.

SMRTbell libraries for the 2-3kb and 3-6kb size-bins were subjected to an additional round of size-selection on the Sage BluePippin to remove trace amounts of small inserts. Primer was annealed and samples were sequenced on the PacBio RS II (Menlo Park) using standard protocols for each respective size-bin. A total of 4 SMRTcells per size-bin were sequenced using P6-C4 chemistry and 4-hour movies.

All the primers and adapter are listed in the **Table S 1**.

### qPCR

#### SMRT-Cappable-seq enrichment

The control and enriched first strand cDNAs were used as templates to determine the enrichment of primary transcript. The level of rRNA and mRNA was examined using qPCR (Luna Universal qPCR Master Mix, New England Biolabs) with primers targeting *E. coli* rrlH, rrlA, hupB,rmf genes and the Gluc gene.

#### Deletion strains

To examine the degree of read-through across the dapA-bamC termination site, the cDNA was synthesized from RNA of wild-type and deletion strain using ProtoScript II Reverse Transcriptase (New England Biolabs) with random primers following the manustructure's instructions. The degree of read-through was measured by comparing the expression level of RNA upstream of TTS (qPCR1) with the level downstream of TTS (qPCR2) (**Figure S6**) using qPCR (Luna Universal qPCR Master Mix, New England Biolabs).

All the qPCR primer sequences are listed in the **Table S 1.**

### Genomic Modifications

To determine the effect of riboswitch-like structure on transcription termination, we deleted a predicted attenuator region of the dapA-bamC terminator (**Figure S6**) from *E. coli* genome (deletion strain) and compared the read-through across this termination site. An *E. coli* B strain C2566 (New England Biolab), referred to in the text as “wild-type,” was used as the parental strain for genomic modification as described [33]. The wild-type and deletion strain were grown in M9 and Rich grown condition late log phase and the RNA was extracted using RNAeasy Mini kit (Qiagen).

### Mapping

The SMRT-seq CCS (circular-consensus) reads were processed using the pacbio_trim.py script. This script contains 2 functions : the filter and poly-function. The filter function removes chimeric reads containing more than one AdaptorL (**Table S 1**) and reads without adaptor L. The remaining reads were trimmed for 3’end polyA sequences and 5’end polyC sequences using the poly-function.

The trimmed reads were mapped to the *E. coli* genome using BLASR (RRID:SCR_000764) (clipping soft). The alignment and all analyses were based on the *E. coli* genome NC_000913.3 and corresponding gene annotation expected for the rRNA analysis.

Additionally, we used adjust_3end.py script on the mapped reads to trim the remaining of polyA tails at the 3’end of the mapped reads.

The RNA-seq Illumina data downloaded from the European Nucleotide Archive (SRR3132588) were trimmed using Trimgalore (http://www.bioinformatics.babraham.ac.uk/projects/trim_galore/) and mapped to the *E. coli* genome NC_000913.3 using BWA (RRID:SCR_010910) version 0.7.5a-r418 [35]. NCBI gene annotation from NC_000913.3 was used to define gene body and the gene expression level was estimated using bedtools (RRID:SCR_006646) multicov version v2.24.0 (-s parameter). For each gene, the number of Illumina reads was compared with the number of PacBio reads overlapping with the gene.

### processed/primary rRNA and protein coding gene classification

Reads were classified based on the overlap with annotated genomic features from *E. coli* U00096.2 annotation (NCBI). Overlap calculations were performed using bedtools intersect version v2.24.0 (-s parameter). To distinguish primary from processed rRNA, the primary known TSS sites for rRNA genes were derived from previous study [4]. Reads starting at the known primary TSS sites for rRNA were classified as primary rRNA, and the remaining reads that overlap with the rRNA genes but do not start at the primary TSS sites were classified as rRNA. The remaining mappable reads were assigned as protein coding genes. The results of this analysis were plotted in **Figure 1B**.

### TSS determination

To identify TSS, we defined T as the number of reads in the same orientation starting at a given position and TPM (tag per million) as T divided by the number of mappable reads multiplying by one million. TPMs were calculated for each position on the genome. We calculated a SMRTratio, which is the ratio of SMRT-Cappable-seq TPM divided by Control TPM. For this analysis, a pseudo-count of 1 has been added to T at each position. Only positions with a SMRTratio above 1 were retained.

Nearby TSS positions in the same orientation within 5 bp distance were clustered together and only the position with the largest number of reads was used as the representative TSS position for the cluster. The score of the TSS corresponds to the sum of T in the cluster.

Only TSSs with scores above 4 were retained in the final set of TSS. 2186 and 1902 TSS were found for the M9 and Rich medium growth condition respectively. TSS were considered as common between growth conditions if their genomic positions are identical and on the same strand. The pearson correlation of TSS between conditions is done by comparing TSS scores between M9 and Rich growth conditions.

The counting and clustering were performed using TSS_analysis.py count and cluster function.

The SMRT-Cappable-seq clustered TSSs under M9 growth condition were compared with the previous Cappable-seq results [4] as described in the Supplementary Text.

### TTS determination

To determine the transcription termination sites (TTS), first the number of transcripts with unique 3' ends was counted for each clustered TSS using TSS_analysis.py count function. To distinguish true TTS from random read end due to incomplete transcription or degradation, we performed a statistical binomial test. More specifically, we calculated the the distance (D in bp from) at the 3' end of a read to its TSS. And if the percent (the number of reads ending at this position divided by the number of reads with the same TSS) was 200 times higher than the probability of random ending at this position (which is 1/D) and there were at least 10 reads, we defined this position as a TTS. The consecutive binomial probabilities were calculated using pbinom function in R, and one-tailed test at the 95% confidence level was used to delimit the statistically significant ends (TTS). The binomial test was performed using the binomial test.R script. Then the nearby TTS positions within 5bp sharing the same TSS were clustered into a TTS cluster. The 5' boundary and 3' boundary of the TTS cluster, and the number of reads that start at the same TSS and end in the TTS cluster were calculated and shown in **Table S2**.

The defined clustered TTSs were compared with the previously experimentally identified termination sites and attenuator sites from Ecocyc (RRID:SCR_002433) [36] as described in the Supplementary Text.

### RNA structure analysis at transcriptional termination sites

RNA sequence 40 bp upstream of the defined TTS was used for predicting secondary structure using RNAstructure [37]. The RNA structure with the lowest free energy was predicted using the Fold function with the following parameters: --loop 30 --maximum 20 --percent 10 --temperature 310.15 --window 3. The RNA structure with free energy lower than −10 was defined as having a termination loop in **Figure 4D**.

### Antiterminator prediction at transcriptional termination sites

RNA sequence 100 bp upstream of the defined TTS was used to predict the antiterminator structure using PASIFIC webserver [32] with parameters optimized for detecting the attenuator. The RNA sequence showing a score above the 0.5 threshold was reported as having an attenuator in **Figure 4D**.

### Motif logo analysis at TTS and TSS

The −30 bp to 10 bp region of the defined TTS and the −45 bp to 5 bp region of the defined TSS under M9 condition was used for motif analysis. Motif logos were generated using the program weblogo3 [38].

### Extended RegulonDB operons

In this study, operons were determined as unique combination of genes included in individual transcripts. Therefore, reads fully covering the same genes were considered to be from the same operon.

The previously annotated operons in RegulonDB were compared with the SMRT-Cappable-seq reads from the combined M9 and Rich dataset. If the read(s) fully covers an annotated operon and an additional gene(s), this annotated operon was extended by SMRT-Cappable-seq. Covering was determined using bedtools intersect version v2.24.0 (-s parameter). The extended RegulonDB operons are shown in **Table S4**.

### SMRT-Cappable-seq operons determination

To determine the transcription unit (TU) based on the SMRT-Cappable-seq result with high confidence, only transcripts with precise TSS (with over 4 reads) were used for this analysis as mentioned above. The start and end of the TU were defined by the TSS and TTS, respectively. For those transcripts without a defined TTS, the longest transcript was used for unit determination. Only the genes that were fully covered by SMRT-Cappable-seq transcripts were annotated into the transcription unit.

Finally the SMRT-Cappable-seq defined operons from both M9 and Rich conditions were compared with the previous annotation in RegulonDB [18] (Supplementary text). The defined SMRT-Cappable-seq operons are shown in **Table S3**.

### Gene context analysis

For each annotated gene, the number of operons that include the gene was calculated, and the gene contexts between M9, Rich condition and RegulonDB (RRID:SCR_003499) were compared.

### Go enrichment analysis

Go enrichment analysis and p.value for genes with additional contexts in Rich medium was performed using the Gene Ontology Consortium [39]. The Bonferroni correction for multiple testing has been applied.

